# Bidirectional regulation between circadian feeding behavior and specialized midgut clocks in *Drosophila*

**DOI:** 10.64898/2026.04.23.720414

**Authors:** Binbin Wu, William W. Ja

## Abstract

Feeding is a fundamental animal behavior. Increasing evidence suggests that the timing of food intake—rather than the amount or quality alone—contributes to maintaining health. Although mistimed eating can reset peripheral clocks and desynchronize them from central pacemakers to affect physiology and metabolism, how peripheral clocks can in turn shape rhythmic feeding behavior is less well understood. Here, we investigated the contribution of peripheral clocks to circadian feeding behavior in *Drosophila* males. Using high-resolution feeding assays combined with complementary analytical approaches that assess both rhythmicity and time-resolved dynamics, we examined the roles of distinct peripheral cell types in feeding regulation. This study reveals the involvement of midgut enteroendocrine cells (EECs) and enterocytes (ECs) in maintaining the stability and strength of feeding rhythms, whereas the fat body clock modulates baseline levels of food intake. Beyond serving as an output behavior of circadian rhythms, feeding also acts as an effective behavioral Zeitgeber that drives molecular clocks. In the absence of the dominant Zeitgeber––light––midgut oscillations decay during prolonged *ad libitum* feeding in constant darkness, whereas feeding/fasting cycles enable the autonomous persistence of clock oscillations in EECs but not in ECs. These findings are suggestive of regulatory feedback between midgut EECs and feeding, highlighting how timed feeding or dietary interventions could influence metabolic health via specialized gut cells.

## Introduction

Circadian rhythms align physiology and behavior with daily environmental cycles to promote survival and fitness^1,2^. These rhythms are coordinated by central pacemakers in the brain that synchronize peripheral oscillators distributed across tissues. Environmental and behavioral cues, such as time-restricted feeding (TRF), can reset peripheral clocks (e.g., in the liver) and uncouple them from the central clock, contributing to metabolic disruption^3,4^. However, it remains unclear whether these peripheral responses are transient or whether they can support sustained feedback between molecular oscillations and feeding rhythms.

Metabolic tissues such as the gut act as endocrine organs that integrate nutritional cues and communicate with the brain through hormone-like signals. In mammals, a prominent example is glucagon-like peptide-1 (GLP-1), a gut-derived peptide hormone that suppresses appetite through the gut-brain axis^5^. These observations raise the possibility that peripheral gut clocks could both respond to and shape feeding behavior.

In *Drosophila*, feeding behavior exhibits a robust circadian pattern, with a preference for food intake during the morning phase^6,7^. Central clock neurons exert broad control over circadian outputs, whereas peripheral clocks are thought to modulate specific behaviors in a tissue- and context-dependent manner. Peripheral clocks have been identified in metabolically active tissues including the fat body, midgut, and glia^8–10^. These oscillators possess the molecular machinery required for self-sustained circadian rhythms, yet their functional contributions differ across tissues and cell types. The adult midgut comprises intestinal stem cells (ISCs), enteroblasts (EBs), enterocytes (ECs), and enteroendocrine cells (EECs)^11^, which show heterogeneous clock gene expression and oscillations^10,12^. TRF improves gene-expression rhythms and metabolic health^13^, and intermittent TRF (iTRF) delays gut aging and extends lifespan in *Drosophila*^14^, suggesting that specific midgut cell types may decode rhythmic feeding cues. However, how distinct peripheral clocks integrate feeding signals and, in turn, help generate or stabilize circadian feeding rhythms remains poorly defined.

Here, we systematically dissect the contributions of fat body and midgut clocks to feeding rhythms and ask whether rhythmic feeding can selectively entrain midgut clocks. To address these questions, we establish and validate a quantitative analytical framework for circadian feeding behavior, examine how peripheral clocks in the fat body and different midgut cell types shape feeding rhythms, and show that rhythmic feeding can selectively entrain midgut clocks, thereby revealing bidirectional regulation between gut clocks and feeding behavior.

## Results

### Detection of rhythmic feeding behavior

To observe circadian rhythms in feeding behavior, we monitored food intake of individually housed *Drosophila* using the ARC system (Figure 1A)^15^. Flies were continuously tracked for 6 days, with 3 days of entrainment under 12-/12-hr light/dark (LD) cycles followed by 3 days in constant darkness (DD) to assess free-running rhythms. Because individual food intake is sparse and highly variable, we characterized circadian feeding rhythms from population-averaged consumption. To validate this detection and analytical strategy, we first performed a pilot experiment using white-eyed Canton-S (*w*CS) flies. These flies exhibited a robust feeding pattern––a prominent peak of consumption in the subjective morning and a trough in the subjective evening––that persisted throughout the free-running period in DD (Figure 1B).

**Figure 1:**
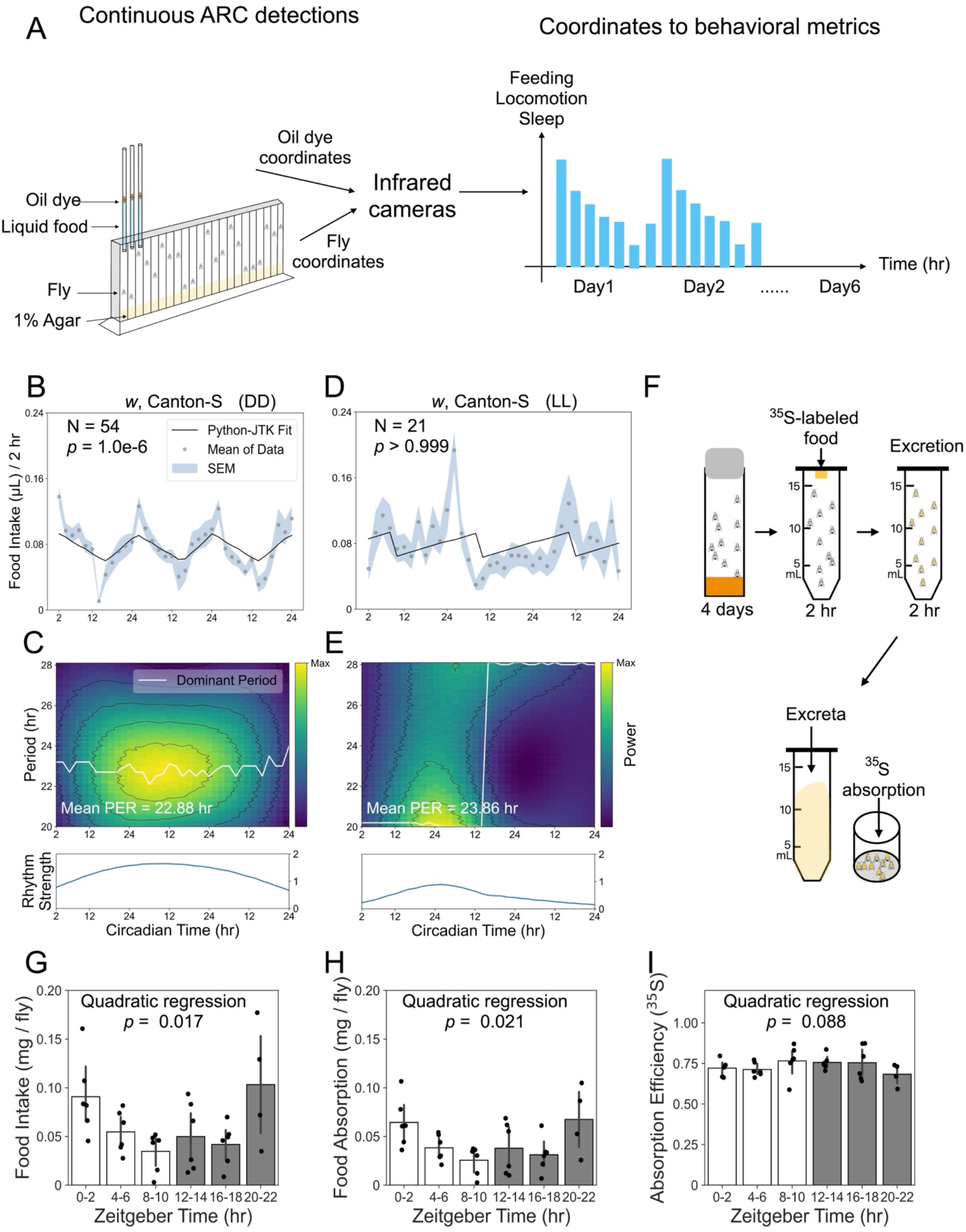
Detection and validation of circadian feeding rhythms in *Drosophila*. (A) Schematic of the ARC-based behavioral measurements (e.g., feeding, sleep, and locomotion), enabling 6 days of continuous tracking from entrainment through the free-running period. Flies are housed individually in chambers and fed liquid food in glass microcapillaries. Feeding is quantified by using infrared cameras to track the meniscus, which is labeled with an oil layer containing an oil-soluble, infrared-absorbing dye. (B) *Drosophila* food intake during the 3-day free-running period (DD1-3) was monitored following LD entrainment. The population-mean food intake was analyzed by Python-JTK. The concordance between the feeding curve and the best-fitting template was assessed by *p* values, with *p* < 0.05 indicating a statistically significant circadian feeding rhythm. (C) CWT analysis characterizes how the period and strength of oscillations change over time. The top panel (heatmap) displays the power of period-specific components that match the feeding signal at each timepoint, while the dominant period (white line) indicates, at each timepoint, the period with the highest power. The bottom panel represents oscillation strength as the globally normalized power of the dominant rhythm over time, enabling direct comparisons between groups. (D) *Drosophila* food intake was monitored under constant light (LL) following LD entrainment. The population-mean food intake during the last 72 hr in LL was analyzed by Python-JTK; *p* > 0.05 indicates no significant circadian rhythmic pattern in feeding. (E) CWT analysis under LL reveals destabilized periods and low oscillation strength over time. (F) Schematic of the ^35^S-labeling method used to measure 2-hr food intake and nutrient absorption over a 24-hr period. At the start of each 2-hr assay period, flies are transferred to tubes containing ^35^S-met-labeled food. After feeding, flies are transferred to empty tubes to collect excreta. After 2 hr, ^35^S is quantified separately from fly bodies and excreta. (G) Food intake was estimated by summing ^35^S retained in the fly bodies and recovered in excreta. Quadratic regression revealed a U-shaped oscillation in food intake over time (N = 5 groups of 10 flies each per condition). (H) Nutrient absorption, estimated by the ^35^S retained in fly bodies, also exhibited a U-shaped oscillation over time. (I) Absorption efficiency, calculated as the ratio of fly body-containing ^35^S to total food intake, remained relatively constant over time. Data in (G-I) are shown as mean ± s.e.m.

To statistically assess endogenous rhythmicity, we applied two complementary analyses implemented in the easyClock software package^16^: Python-JTK and continuous wavelet transform (CWT). Python-JTK tests whether the feeding time series follows a circadian (∼24-hr) rhythm by comparing the observed pattern to reference rhythmic waveforms, yielding a *p*-value that indicates whether feeding is significantly rhythmic over the free-running period (Figure 1B). In contrast, CWT analysis captures how rhythmic feeding patterns evolve over time. Rather than assuming a stationary rhythm, CWT decomposes the feeding signal into time-varying oscillations, allowing simultaneous assessment of period and rhythm strength. In the CWT heatmap, color represents wavelet power (rhythm strength) across different periods, and the dominant period (white line) indicates, at each timepoint, the period with the highest power, reflecting period stability (Figure 1C, upper). To compare rhythm strength between groups, we normalized wavelet power to the maximum mean power of the dominant rhythms across groups, producing a rhythm-strength curve (Figure 1C, lower). Conceptually, Python-JTK determines whether a rhythm is present, whereas CWT evaluates the robustness and stability of that rhythm. A significant Python-JTK result with weak CWT metrics (e.g., reduced rhythm strength or drifting periods) indicates that the rhythmic feeding pattern is present but not well sustained. Conversely, strong and stable oscillations identified by CWT are typically accompanied by highly significant Python-JTK results, reflecting that the rhythmic feeding pattern is detectable and well sustained. Together, these two approaches provide a detailed and dynamic characterization of feeding behavior.

In chronobiology, constant light exposure disrupts circadian rhythms through promoting CRYPTOCHROME (CRY)-mediated TIMELESS (TIM) degradation^1,17^. To test whether ARC-detected feeding rhythms are similarly abolished, we exposed flies to constant light following LD entrainment. As anticipated, no significant circadian rhythmic pattern was detected in population-averaged feeding (Figure 1D), and CWT analysis revealed destabilized feeding periods and dampened oscillation strength (Figure 1E).

To further clarify whether ARC-quantified feeding oscillations accurately reflect variations in food intake and nutrient absorption, we used a radioisotope-labeling assay to quantify 2-hr food intake and estimate nutrient absorption at different timepoints (Figure 1F). Both food intake and absolute absorption exhibited a U-shaped curve over time (Figure 1G–H), closely matching the ARC-quantified feeding pattern (Figure 1B). In contrast, absorption efficiency, calculated as the ratio of absorbed radiotracer to total intake, remained relatively constant over time (Figure 1I). Spearman correlation analysis further revealed a strong positive association between food intake and absorption (*r* = 0.97; *p* = 8e-22), whereas absorption efficiency and food intake were negatively correlated (*r* = -0.51; *p* = 0.002). Together, these results validate our strategy for detecting endogenous circadian rhythms in feeding behavior.

### Global disruption of core clock genes eliminates circadian feeding behavior

Compared with changes in photoperiod, genetic perturbations of core clock genes provide a more flexible approach for investigating endogenous feeding rhythms. We first examined *Clk*^jrk^, a mutant allele of the core clock gene, *Clock* (*Clk*), which is known to regulate circadian rhythms, including rhythmic feeding^6,7,18^. As expected, *Clk*^jrk^ mutants failed to maintain a rhythmic pattern in feeding behavior during the free-running period compared to the iso^31^ control (Figure 2A, C). Although *Clk*^jrk^ flies maintained stable circadian periods, their feeding oscillations showed markedly reduced strength (Figure 2B, D), resulting in weakened rhythmic output that failed template-based rhythmicity detection. This dissociation highlights that rhythmicity detection and oscillation strength can diverge.

**Figure 2:**
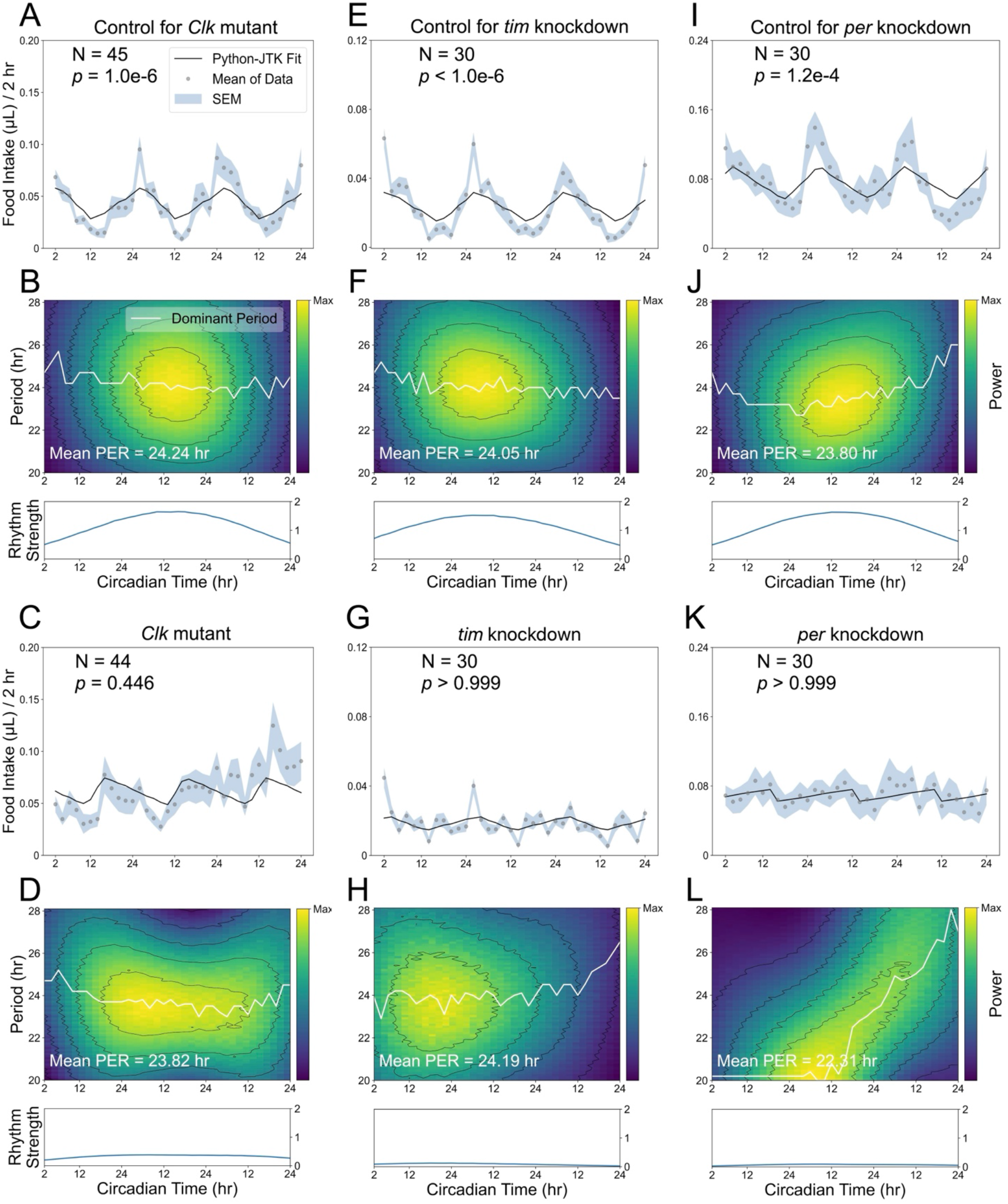
Global disruption of core clock genes abolishes circadian feeding rhythms. All flies experienced 3 days of LD followed by 3 days of DD in the ARC system, and feeding rhythmicity during the free-running period was analyzed using Python-JTK and CWT. (A, C) Population-mean food intake during DD1-3 for iso^31^ controls (A), in which Python-JTK detects a significant circadian feeding rhythm, and *Clk*^jrk^ mutants (C), in which no significant rhythm is detected. (B, D) CWT analysis shows that *Clk*^jrk^ mutants (D) exhibit reduced oscillation strength compared to iso^31^ controls (B). (E, G) Population-mean food intake for flies in which *acp* (control, E) or *tim* (G) was targeted by *tim*-GAL4-driven CRISPR-Cas9. Python-JTK detects a significant feeding rhythm in *acp*-targeted controls (E), but not in *tim* knockouts (G). (F, H) CWT analysis indicates that *tim*-knockout flies (H) show substantially lower oscillation strength than their controls (F). (I, K) Population-mean food intake for *acp*-targeted controls (I) and *per* knockouts (K) through *tim*-GAL4-driven CRISPR-Cas9, with Python-JTK detecting a significant circadian feeding rhythm only in controls. (J, L) CWT analysis indicates that *per*-knockout flies (L) lose period stability and show reduction in oscillation strength over time.

We further used a GAL4-driven CRISPR-Cas9 approach to globally disrupt the core clock genes *timeless* (*tim*) or *period* (*per*). This tool uses the GAL4/UAS binary system to drive Cas9 and single guide RNA (sgRNA) expression *in vivo*, enabling cell type-specific gene disruption^19^. As a negative control, targeting *accessory gland protein 98AB* (*acp*) in *tim*-expressing cells had no effect on feeding rhythms, whereas knocking out *tim* or *per* in these cells abolished circadian feeding patterns (Figure 2E-L).

To screen peripheral clocks related to feeding, we expressed a dominant-negative form of *Clk* (*Clk*.Δ) that disrupts CLK/CYC complex formation. *Clk*.Δ encodes a truncated protein that competitively binds intracellular CYC but lacks the basic region required for activating downstream gene expression^20^. Because broad *Clk*.Δ expression during development compromises *Drosophila* viability, we introduced temperature-sensitive GAL80 (GAL80^ts^) to temporally control genetic manipulations. To test the feasibility of this approach at the permissive (22 °C) and restrictive (29 °C) temperatures of GAL80^ts^, we combined *tub*-GAL80^ts^ with *Clk*.Δ expression to interfere with CLK/CYC in *tim*-expressing cells specifically in adulthood. Induction of *Clk*.Δ expression at 29 °C in adults abolished rhythmic feeding patterns in the experimental group, whereas control genotypes maintained stable, robust feeding rhythms (Figure 3A-F). At 22 °C, feeding rhythms in both control and experimental groups remained intact and robust (Figure 3G-L). These results confirm that our GAL80^ts^-based strategy is suitable for characterizing feeding rhythms at different temperatures for clock-disruption studies.

**Figure 3:**
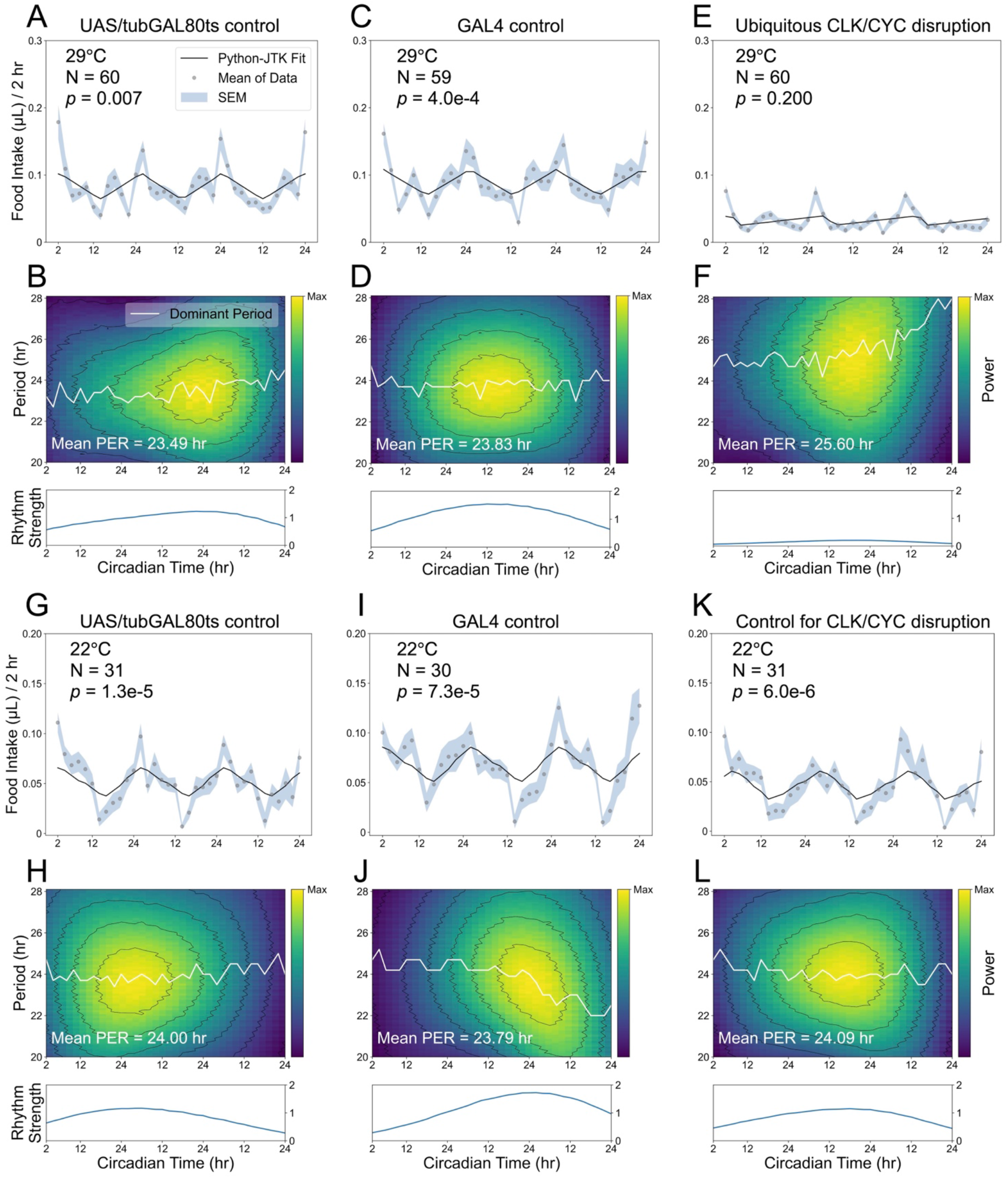
Adult-specific disruption of *Clk* function abolishes circadian feeding rhythms. All flies experienced 3 days of LD followed by 3 days of DD in the ARC system, and feeding rhythmicity during the free-running period was analyzed using Python-JTK and CWT. (A-F) At 29 °C (restrictive temperature), *tim*-GAL4 drives dominant-negative *Clk* (*Clk*.Δ) expression in *tim*-expressing cells (E, F: *tim* > *Clk*.Δ, *tub*-GAL80^ts^), resulting in arrhythmic feeding patterns, destabilized periods, and weakened oscillations compared with controls (A, B: *w*CS; UAS-*Clk*.Δ, *tub*-GAL80^ts^/+; C, D: *w*CS; *tim*-GAL4/+). (G-L) At 22 °C (permissive temperature), GAL80^ts^ inhibits GAL4 activity, and all genotypes display robust circadian feeding rhythms, with stable periods and relative high oscillation strength.

### Identifying peripheral clocks associated with feeding

To identify peripheral clocks that regulate rhythmic feeding, we first investigated the fat body due to its well-established role in metabolism. Disrupting fat body clocks preserved but weakened feeding rhythms, with lower oscillation strength over time (Figure 4A-D). Expressing *Clk*.Δ in the fat body also significantly reduced daily food consumption (Figure 4E). We verified that this fat body driver is strongly expressed in the fat body (Figure 4F), but also shows off-target expression in midgut ECs, including midgut R1 and a portion of anterior R2 (Figure 4G). Although it is not expressed in clock neurons (Figure 4H), this partial perturbation of midgut clocks may confound interpretations of the role of the fat body in feeding regulation.

**Figure 4:**
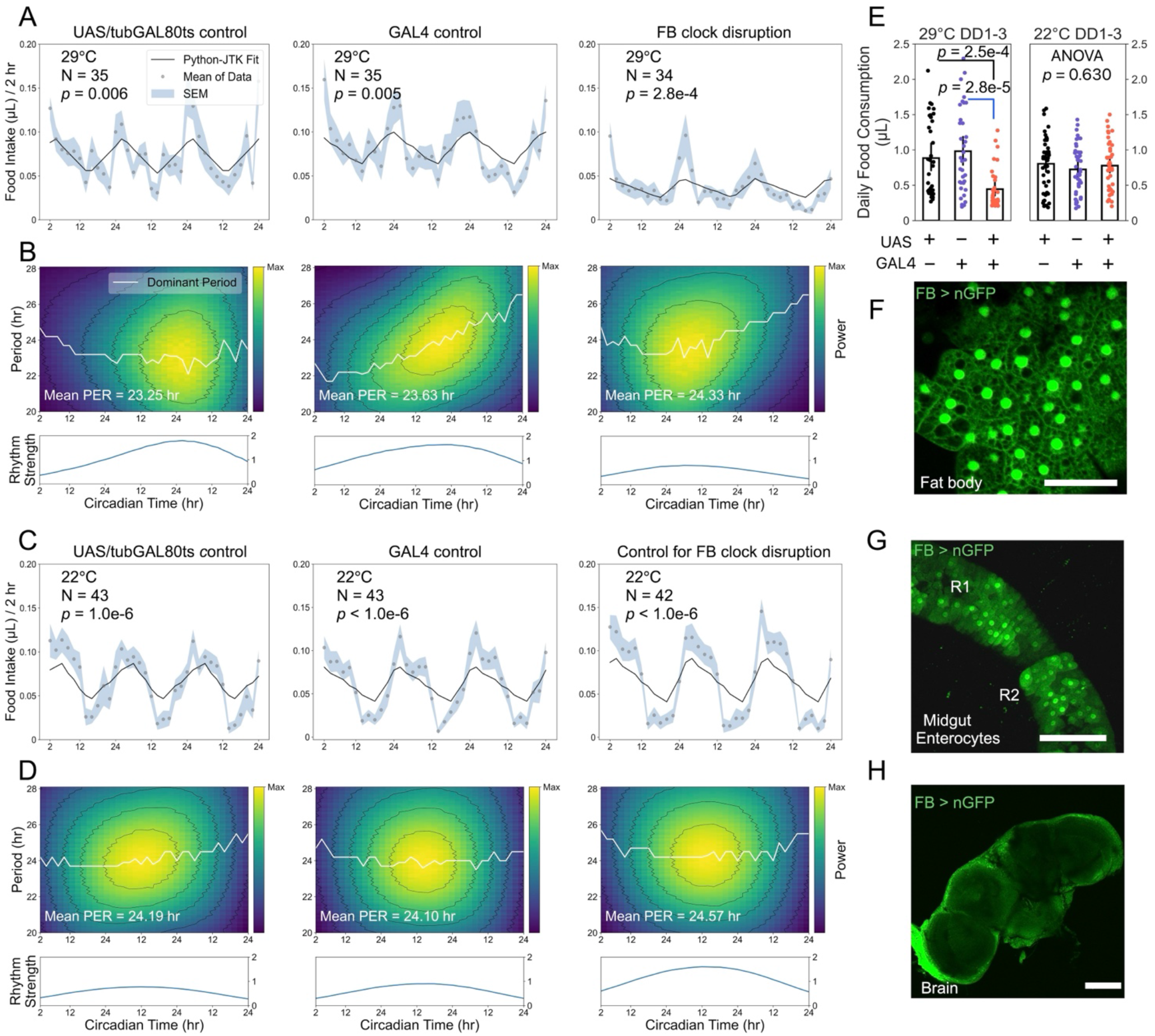
Fat body clock disruption weakens feeding rhythms and reduces food intake. All flies experienced 3 days of LD followed by 3 days of DD in the ARC system, and feeding rhythmicity during the free-running period was analyzed using Python-JTK and CWT. (A, B) At 29 °C, fat body driver-mediated *Clk*.Δ expression does not abolish feeding rhythmicity but reduces oscillation strength. (C, D) At 22 °C, all genotypes remain circadian rhythmic, with stable periods and robust feeding oscillations. (E) Fat body driver-mediated *Clk*.Δ expression significantly reduces daily food consumption. Multiple comparisons were conducted using ANOVA (Kruskal-Wallis test) followed by Dunn’s post-hoc test. (F-H) The fat body driver shows robust expressions in fat body (F), mild expression in enterocytes of anterior midgut regions (G), and no evident expression in the brain (H). Scale bars: 50 μm (F) and 100 μm (G, H). Genotypes: *w*CS; UAS-*Clk*.Δ, *tub*-GAL80^ts^/+ (UAS/tubGAL80ts control); *w*CS; FB-GAL4/+ (GAL4 control); FB > UAS-*Clk*.Δ, *tub*-GAL80^ts^ (FB clock disruption).

To further investigate whether midgut ECs contribute to the observed effects, we used the *Myo1A-*GAL4 driver, which is broadly expressed in midgut ECs (Figure 5A-B). Driving *Clk*.Δ expression in ECs during adulthood did not abolish feeding rhythms (Figure 5C), but reduced their robustness and temporal precision, leading to weaker and lengthened oscillations over time (Figure 5D). However, *Myo1A-*GAL4 also exhibits weak expression in the brain (Figure 5E), implying extra-intestinal activity. To exclude neuronal contributions, we introduced *nsyb-*GAL80 to suppress GAL4 activity in neurons (hereafter *Myo1A-*GAL4^GAL80^). Compared with the original driver, *Myo1A-*GAL4^GAL80^ efficiently inhibited GAL4 activity in the brain (Figure 5F). Driving *Clk*.Δ expression using *Myo1A-*GAL4^GAL80^ produced weak but stable feeding rhythms (Figure 5G-H). Importantly, disrupting clocks in midgut ECs using either *Myo1A-*GAL4 or *Myo1A-*GAL4^GAL80^ did not reduce food consumption (Figure 5I-J), supporting a role for fat body clocks in regulating overall food intake, whereas EC clocks primarily influence the strength of circadian feeding rhythms.

**Figure 5:**
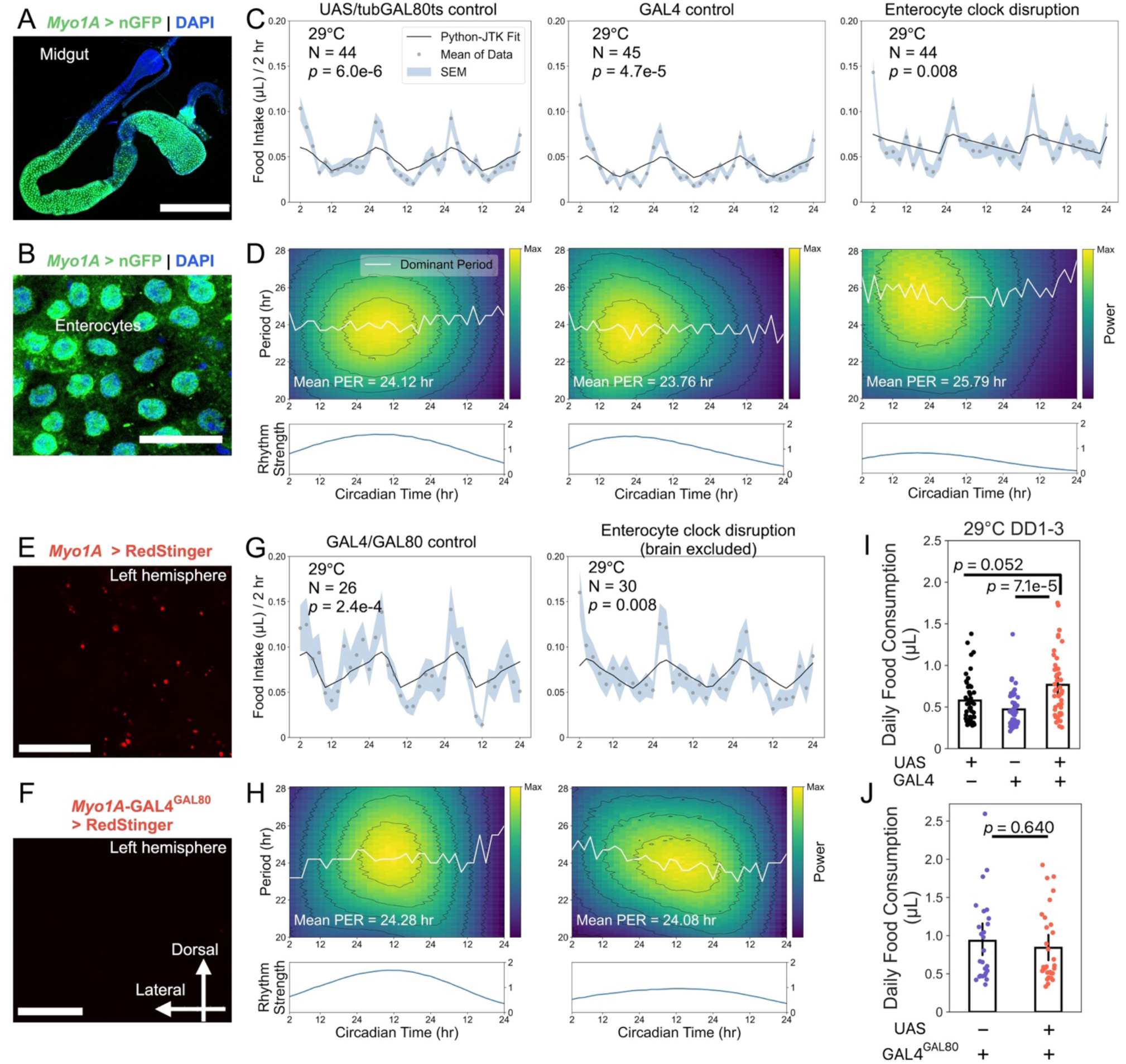
Midgut enterocyte clocks support feeding rhythm strength without altering total food intake. All flies experienced 3 days of LD followed by 3 days of DD in the ARC system, and feeding rhythmicity during the free-running period was analyzed using Python-JTK and CWT. (A) *Myo1A-*GAL4 drives nuclear GFP (nGFP) expression in the midgut. (B) *Myo1A-*GAL4-driven nGFP is confined to enterocytes. Blue: DAPI; green: nGFP (A, B). (C, D) At 29 °C, *Myo1A*-GAL4-driven *Clk*.Δ expression in ECs leads to lengthened periods and dampened oscillation strength. (E) *Myo1A*-GAL4 driving the RFP variant RedStinger (red) reveals mild expression in the brain. (F) Introduction of *nsyb*-GAL80 suppresses GAL4 activity in neurons (*Myo1A*-GAL4^GAL80^), eliminating brain expression. (G, H) At 29 °C, expressing *Clk*.Δ in enterocytes but not in neurons produces weak but stable feeding rhythms with reduced oscillation strength. (I, J) Disrupting the enterocyte clock using *Myo1A*-GAL4 or *Myo1A*-GAL4^GAL80^ does not reduce daily food consumption. Data are presented as mean ± s.e.m. Group comparisons were performed using one-way ANOVA (Kruskal-Wallis test) followed by Dunn’s post-hoc test (I) or the Mann-Whitney U test (J). Scale bars: 400 µm (A), 30 µm (B), 100 µm (E, F). Genotypes: *w*CS; UAS-*Clk*.Δ, *tub*-GAL80^ts^/+ (UAS/tubGAL80ts control); *w*CS; *Myo1A*-GAL4/+ (GAL4 control); *Myo1* > UAS-*Clk*.Δ, *tub*-GAL80^ts^ (enterocyte clock disruption); *Myo1A*-GAL4^GAL80^ (GAL4/GAL80 control); *Myo1A*-GAL4^GAL80^ > UAS-*Clk*.Δ, *tub*-GAL80^ts^ (enterocyte clock disruption with brain excluded).

In mammals, peptides released from EECs, such as GLP-1, potently suppress appetite and control body weight through the gut-brain axis^5^. GLP-1 secretion displays circadian oscillations in mammals^21,22^, suggesting that EEC clocks may also contribute to the regulation of feeding rhythms. In *Drosophila* single-cell RNA sequencing indicates that EECs express very low mRNA levels of core clock genes such as *tim, cwo, pdp1*, and *vri*^10^, and the physiological function, if any, of CLK/CYC-expressing EECs in feeding regulation remains unclear. The interpretation is further complicated by off-target brain expression shown by commonly used EEC drivers. We avoided such non-specific manipulations by using a previously identified split-GAL4 pair (R20C06-*p65*.AD ⋂R33A12-GAL4.DBD)^23^ to investigate the involvement of EECs in circadian feeding while bypassing brain contributions. Confocal imaging confirmed strong and broad expression in midgut EECs with minimal brain expression (Figure 6A-B). Disrupting clocks in these EECs with *Clk*.Δ expression resulted in an initially lengthened but progressively shortened circadian period with reduced oscillation strength, indicating weakened and destabilized rhythmic feeding (Figure 6C-D). By contrast, in less differentiated midgut cell populations, including ISCs and EBs (Figure 6E-F), adult-specific *Clk*.Δ expression driven by an RU486-inducible GeneSwitch system did not disrupt circadian feeding rhythms (Figure 6G).

**Figure 6:**
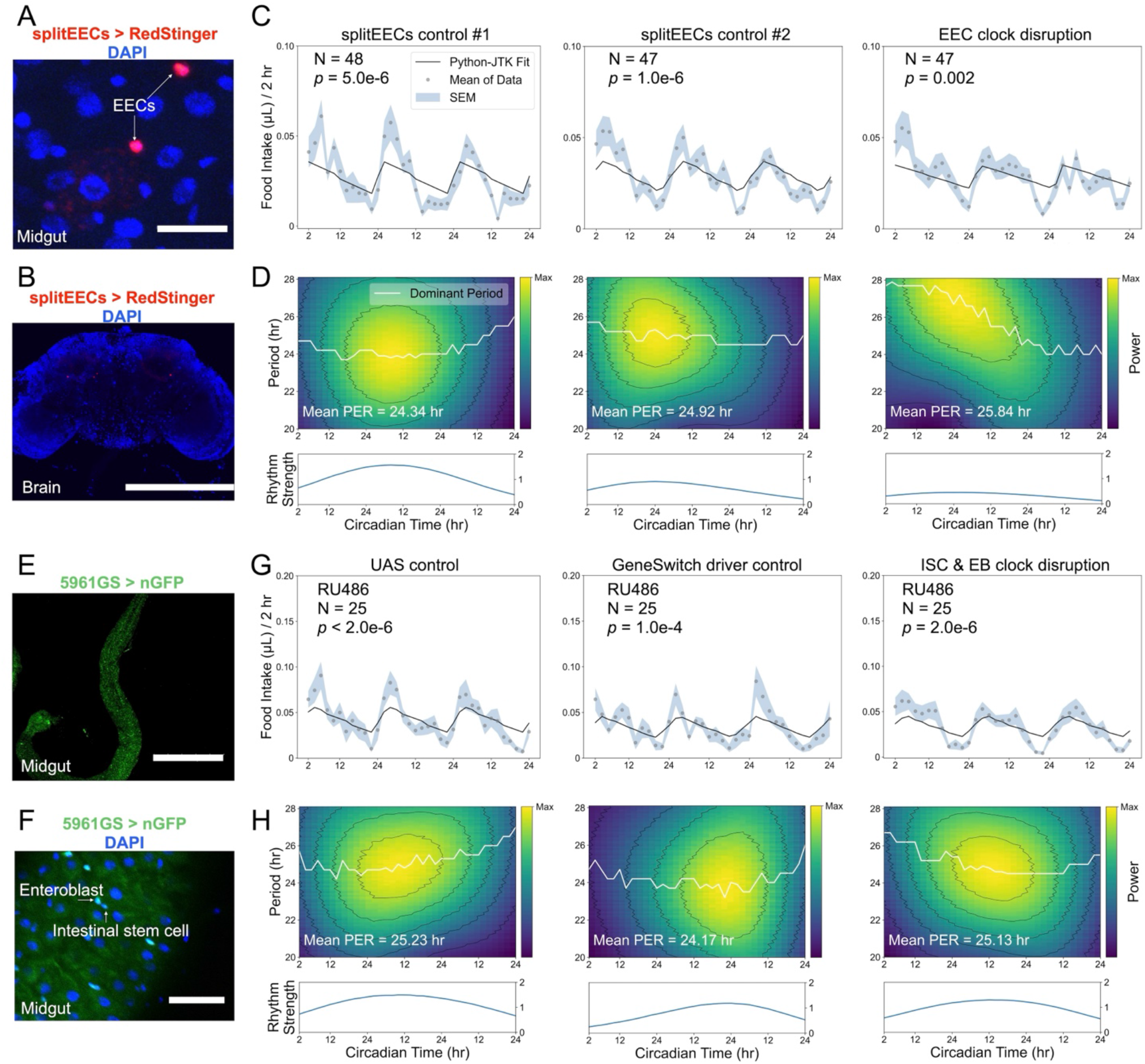
Midgut enteroendocrine clocks stabilize circadian feeding rhythms. All flies experienced three days of LD followed by three days of DD in the ARC system, and feeding rhythmicity during the free-running period was analyzed using Python-JTK and CWT. (A, B) The split-GAL4 drivers together (splitEECs: R20C06-p65.AD ∩ R33A12-GAL4.DBD) drive RedStinger expression in midgut enteroendocrine cells (EECs, A) but not in the brain (B). Blue: DAPI; red: RedStinger. Scale bars: 20 µm (A) and 300 µm (B). (C, D) Expressing *Clk*.Δ in midgut EECs destabilizes circadian periods and reduces oscillation strength, indicating weakened and destabilized feeding rhythms. (E, F) The GeneSwitch driver 5961GS induces nGFP expression in progenitor cells (intestinal stem cells, ISCs, and enteroblasts, EBs) upon administration of RU486. Blue: DAPI; green: nGFP. Scale bars: 400 µm (E) and 30 µm (F). (G, H) Adult-specific disruption of CLK/CYC in midgut progenitor cells (ISCs/EBs) does not affect the maintenance of circadian feeding rhythms. Genotypes are as follows: R20C06-*p*65.AD > *Clk*.Δ (splitEECs control #1); R33A12-GAL4-DBD > *Clk*.Δ (splitEECs control #2); R20C06-*p*65.AD ∩ R33A12-GAL4.DBD > *Clk*.Δ (EEC clock disruption); *w*CS; UAS-*Clk*.Δ/+ (UAS control); *w*CS; 5961GS/+ (GeneSwitch driver control); 5961GS > *Clk*.Δ (ISC & EB clock disruption).

### Rhythmic feeding selectively entrains midgut clocks

Given the contributions of ECs and EECs to feeding rhythms, we re-evaluated molecular clock oscillations within the midgut using the reporter, Clock[*per*]. Clock[*per*] expresses a destabilized nuclear GFP under the control of a minimal *per* promoter that is activated by CLK/CYC^12^. Thus, Clock[*per*] GFP intensity provides a time-resolved readout of nuclear CLK/CYC-driven transcriptional activity at the *per* promoter.

The adult *Drosophila* midgut is anatomically divided into five regions (R1-R5) from anterior to posterior, each with distinct morphologies, functions, and transcriptomes, while regions R2 and R4 comprise most of the midgut^24^. Consistent with previous studies^10,12^, we observed robust CLK/CYC oscillations in most regions of midgut ECs under LD (Figure 7A), which were abolished by exposing flies to constant light (Figure 7B). Prior work indicates that this synchronization occurs cell-autonomously in CRY(+) peripheral tissues, including intestinal cells^25^.

**Figure 7:**
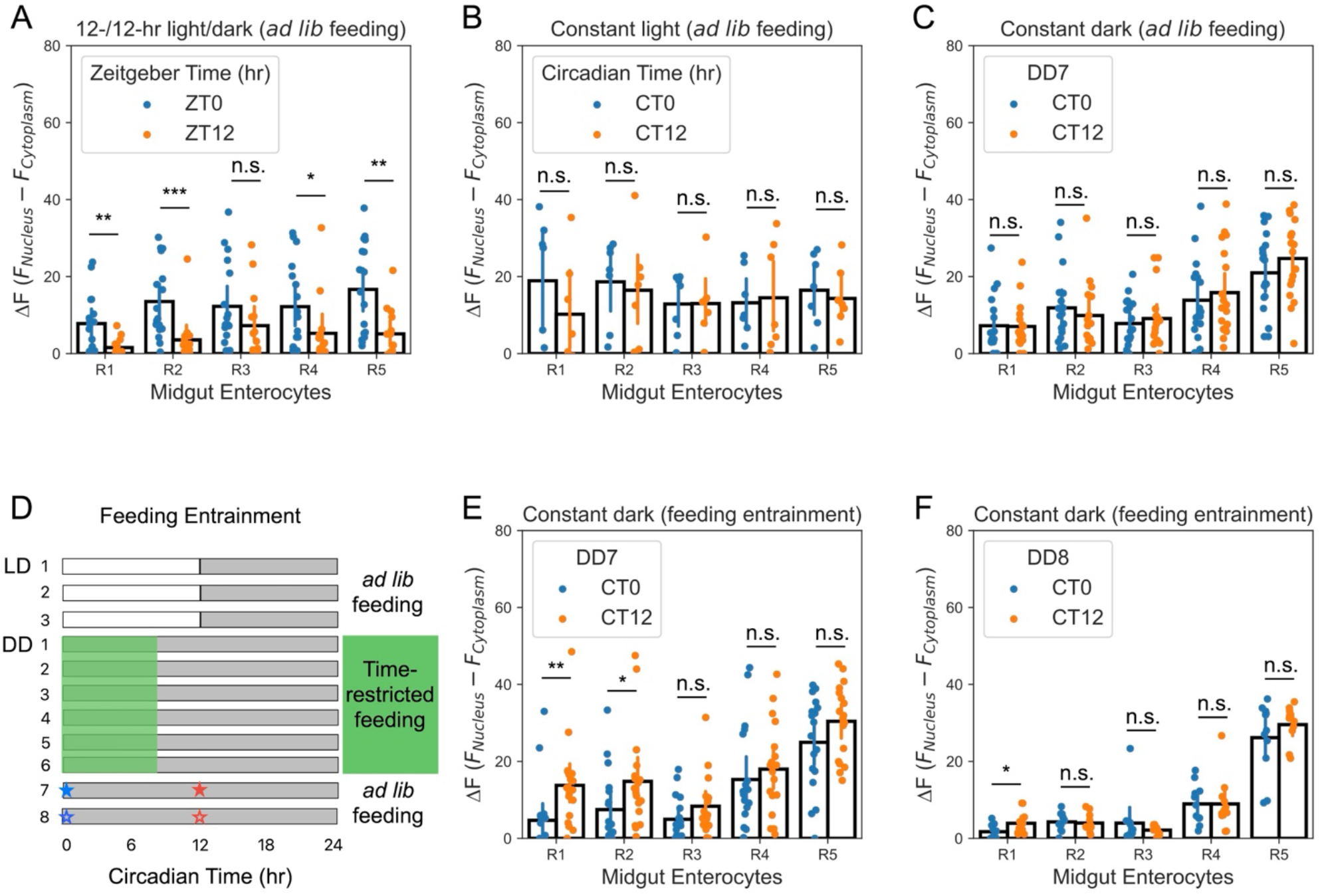
Midgut enterocyte clocks are not stably entrained by feeding/fasting cycles. (A) CLK/CYC activity in enterocytes (EC_s_) from midgut regions R1-R5 under LD. CLK/CYC shows high expression at ZT0 and low expression at ZT12 (N = 14-17). (B) Constant light eliminates enterocyte CLK/CYC oscillations in midgut (N = 7). (C) After seven days of *ad libitum* feeding under constant darkness (DD), flies lose enterocyte CLK/CYC oscillations (N = 18). (D) Schematic of the feeding/fasting entrainment protocol under DD. Flies underwent six cycles of 8-hr feeding and 16-hr fasting in the absence of light, followed by a free-running, *ad libitum* fed period. The markers (★*/*☆) represent the sampling timepoints in (E, ★) and (F, ☆). (E, F) CLK/CYC oscillations in R1-R2 ECs are phase-inverted (“flipped”) after six cycles of feeding/fasting entrainment (E; N = 18) but are largely lost after a subsequent 24-hr free-running period (F; N = 11), with only a residual, but significant, signal in R1. CLK/CYC dynamics were quantified as ΔF = F_nucleus_ – F_cytoplasm_(nuclear GFP – cytoplasmic GFP) using the reporter Clock[*per*]. Bars show mean ± s.e.m. Group comparisons were conducted using the Mann-Whitney U test. *, *p* < 0.05; **, *p* < 0.01; ***, *p* < 0.001; n.s., *p* > 0.05.

We next asked whether the enterocyte clock could be entrained by feeding cues. Under DD with continuous access to food (*ad libitum* feeding) for 7 days (including the test day), enterocyte Clock[*per*] oscillations were lost (Figure 7C), consistent with previous findings that the EC oscillations are not sustained beyond 5 days in DD^9^. To test the effect of rhythmic feeding, Clock[*per*] adults were raised under a feeding/fasting regimen in the absence of light (Figure 7D). The 8-hr feeding/16-hr fasting regimen was chosen based on prior studies of time-restricted feeding, which demonstrate that limiting food intake to 8 hr per day improves metabolic health^26,27^. Surprisingly, consecutive feeding/fasting cycles appeared to invert, rather than synchronize, CLK/CYC oscillations in R1-R2 ECs (Figure 7E), in contrast to their original phase under LD/*ad libitum* conditions (Figure 7A). However, starvation can induce autophagy^14^, and enhanced delivery of GFP to acidic autolysosomes is known to reduce GFP fluorescence in *Drosophila* and other systems^28,29^, potentially decreasing Clock[*per*] signal independently of changes in clock activity. To address this, we re-examined these flies after a 24-hr *ad libitum* feeding period (a free-running period) following feeding/fasting entrainment. Under these conditions, flies no longer showed the “flipped” oscillation pattern in ECs, except for a residual signal in R1 (Figure 7F). Thus, in the absence of light, the enterocyte clock does not show stable entrainment by feeding/fasting cues.

We next investigated the presence of an enteroendocrine clock and its potential entrainment by rhythmic feeding. Confocal imaging showed that a subset of R20C06-*p65*.AD ⋂R33A12-GAL4.DBD-labelled EECs exhibited weak Clock[*per*] GFP signals compared to surrounding ECs at ZT0 (Figure 8A). Because individual EEC Clock[*per*] signals were faint, we assessed EEC clock activity by visually counting Clock[*per*]-positive EECs, defined as EEC nuclei with GFP clearly above the local epithelial background. We then interpreted changes in the number of Clock[*per*]-positive EECs as a semi-quantitative proxy for EEC clock activity. Under LD cycles, the number of CLK(+)-EECs showed a clear oscillation between ZT0 and ZT12 (Figure 8A-C), indicating clock activity in approximately 20 EECs. This oscillation was lost under DD cycles with *ad libitum* feeding (Figure 8D-E). Remarkably, a consistent CLK/CYC oscillation in ∼20 EECs was restored after a free-running period following feeding/fasting entrainment (Figure 8F), indicating that the enteroendocrine clock can be entrained by feeding/fasting cues and that these oscillations are maintained even after feeding cues are removed.

**Figure 8:**
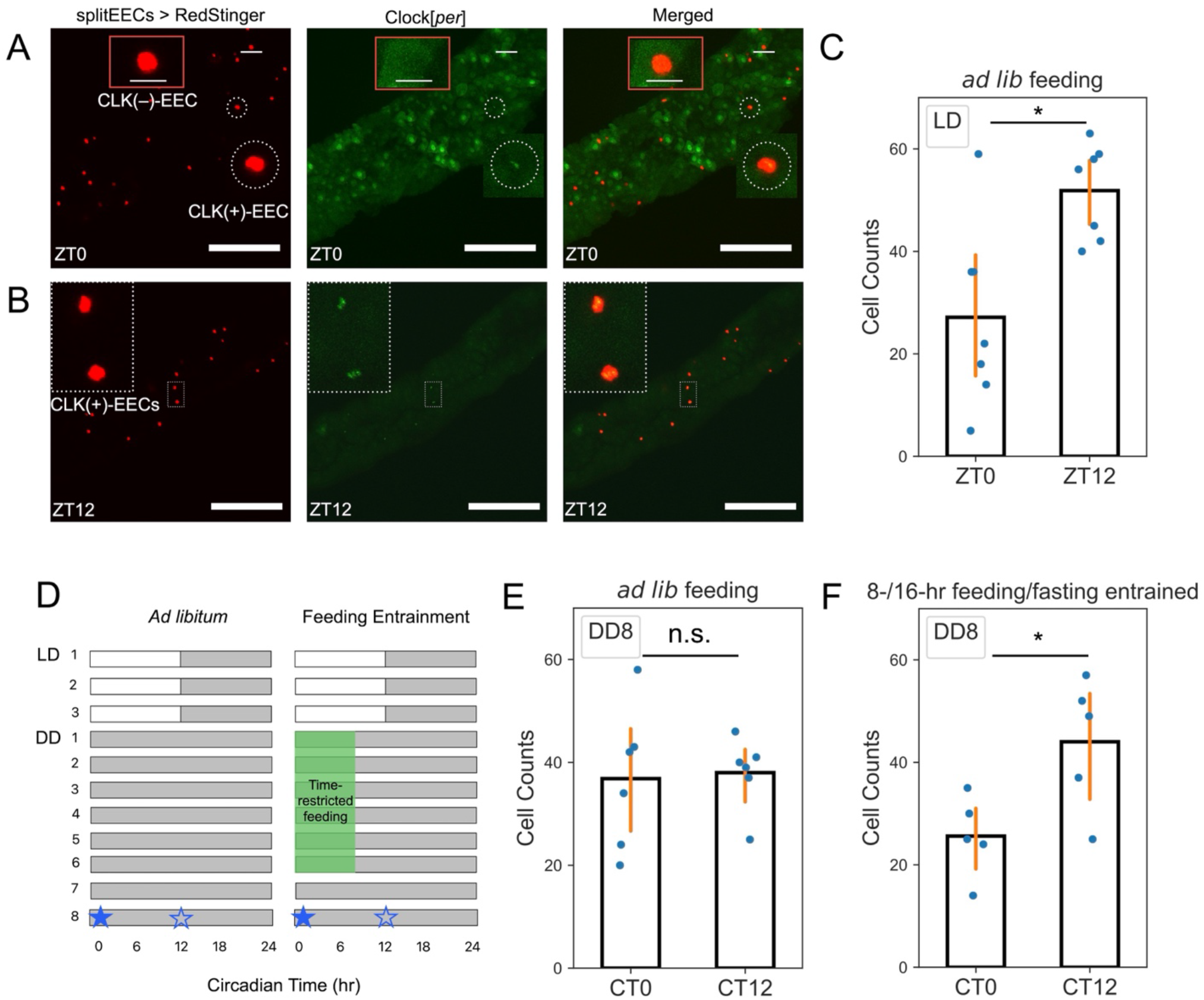
Feeding/fasting cycles entrain and sustain CLK/CYC oscillations in midgut enteroendocrine cells. (A, B) Co-localization of midgut EECs (red, R20C06-*p65*.AD ⋂R33A12-GAL4.DBD splitEECs > RedStinger) with the CLK/CYC reporter Clock[*per*] (green) at ZT0 (A) and ZT12 (B). At ZT0, a CLK/CYC-negative EEC [CLK(–) EEC] is indicated by a red box, and a CLK/CYC-positive EEC [CLK(+) EEC] is circled by white dashed lines; CLK(+) EECs are also visible at ZT12. Scale bars: 100 µm. (C) Number of CLK(+) EECs per midgut at ZT0 and ZT12 under LD (N = 6-7). (D) Schematic of feeding/fasting entrainment protocol under constant darkness (DD). Flies experienced six cycles of 8-hr feeding and 16-hr fasting in the absence of light, followed by a free-running period with *ad libitum* feeding. Control flies were maintained with continuous food access throughout the experiment. Markers (★*/*☆) indicate sampling at CT0 (★) and CT12 (☆) for panels (E) and (F). (E, F) Under DD with seven days of *ad libitum* feeding, EEC CLK/CYC oscillations are lost (E). In contrast, flies subjected to feeding/fasting entrainment exhibit a robust oscillation in the number of CLK(+) EECs that persists after a 24-hr free-running period (F). N = 5-6. Bars show mean ± s.e.m. Group comparisons were conducted using the Mann-Whitney U test. *, *p* < 0.05; n.s., *p* > 0.05.

These results collectively indicate that rhythmic feeding/fasting cycles entrain molecular clocks in EECs but not ECs. Therefore, rather than acting through central oscillators in the brain to synchronize all peripheral clocks, feeding cues appear to act selectively on EECs, which may in turn relay time-of-day signals to the brain or other tissues.

## Discussion

In this study, we dissected how peripheral clocks in the fat body and midgut contribute to circadian feeding behavior, revealing distinct and complementary roles for these tissues. Our results indicate that EEC clocks play a non-redundant role in maintaining stable, robust feeding rhythms and for sustaining clock oscillations in response to rhythmic feeding, whereas EC clocks primarily support the strength of feeding rhythms. In both cases, disrupting midgut clocks in ECs or EECs alters rhythm quality without markedly changing total consumption. Taking advantage of direct measurements of food intake in the ARC, we further show that manipulating fat body clocks reduces baseline consumption, consistent with a role for fat body timekeeping in setting overall energy balance rather than generating feeding rhythms *per se*. Given the off-target expression of our fat body driver in anterior ECs, we conservatively refrain from assigning a pacemaker role to the fat body and instead propose that reduced baseline intake, together with driver-dependent midgut perturbations, may contribute to prior discrepancies regarding fat body control of feeding rhythms across different studies^6,7,30,31^.

Although both EECs and ECs harbor clock machinery, their differential entrainability to feeding/fasting cues points to functional specialization within the midgut. ECs exhibit robust CLK/CYC oscillations under light-dark cycles yet fail to maintain or stably re-entrain these oscillations under prolonged constant darkness with rhythmic feeding, suggesting that they lack sufficient sensory or signaling mechanisms to interpret feeding-related Zeitgeber signals. By contrast, a small subset of EECs shows clock oscillations that are lost during *ad libitum* feeding but restored and self-sustained after feeding/fasting entrainment, consistent with a role for EEC clocks as feedable and persistent oscillators. Notably, Clock[*per*] signal in ECs is higher at ZT0 than at ZT12, whereas EEC Clock[*per*] signal shows the opposite pattern (higher at ZT12 than at ZT0), consistent with cell-type-specific circadian phase variation observed from *Drosophila* to mammals^32–35^. This divergence may reflect differential responses to distinct primary cues or adaptation to cell-type-specific functions. EECs express specialized nutrient sensors^36,37^ and secrete peptide hormones such as Neuropeptide F (NPF) and Allatostatin C (AstC)^36,38^, which influence systemic metabolism and feeding, raising the possibility that circadian regulation of these pathways could relay timing information from gut clocks to the brain and other tissues. These mechanistic links remain to be tested directly, for example by manipulating specific EEC peptides or receptors in a clock-dependent manner.

More broadly, our findings support the view that peripheral clocks do not simply follow instructions from the central clock but can shape circadian output in a tissue-specific and behaviorally relevant manner. While the central brain clock is essential for coordinating organism-wide timing, our results suggest that midgut EECs can act as partially autonomous oscillators that both respond to and stabilize rhythmic feeding. In this sense, circadian feeding emerges from a distributed system in which central and peripheral clocks interact with behavioral feedback rather than a strictly top-down hierarchy. This framework may help explain why timed feeding or dietary interventions can have potent effects on metabolic health: by selectively engaging gut clocks in cells analogous to EECs, such regimens could reinforce or restore local clock function even when systemic timing cues are perturbed.

## Materials and methods

### *Drosophila* stocks and maintenance

Flies were reared under 12-/12-hr light/dark cycles at 25 °C. CRISPR-Cas9-related UAS lines, 5961GS, iso^31^, and *Clk*^jrk^ were obtained from the laboratory of Mimi Shirasu-Hiza; *tim-*GAL4 (#7126), UAS-nGFP (#4775), UAS-RedStinger (#8547), *nsyb*-GAL80 (#92153), and split EECs drivers (#70585 and #68537) were obtained from the Bloomington *Drosophila* Stock Center; Clock[*per*] was obtained from the laboratory of Phillip Karpowicz. Fly lines were outcrossed to *w*CS for approximately 10 generations to standardize genetic background, except *Clk*^jrk^, which was maintained in the iso^31^ background. For CRISPR-Cas9 experiments, controls carried a construct targeting an unrelated gene (*acp*), whereas experimental flies carried constructs targeting the gene of interest.

### Inducible genetic manipulations

For thermogenetic experiments using GAL80^ts^, flies were raised at 18 °C during development. Behavioral experiments were conducted at 22 °C for inhibiting GAL4-induced expression or at 29 °C for inducing GAL4-activating effects.

For drug-inducible GeneSwitch studies, the 5961GS driver was induced by feeding 186 μM (80 μg/mL) RU486. Stock RU486 (40 mg/mL) in ethanol was diluted into liquid food to achieve the desired concentration (186 µM). Flies were provided with food containing RU486 or vehicle (ethanol) from the acclimation day to the end of the experiment (a total of 6 days). In the GFP labelling experiments, flies were fed for RU486 for 2-3 days.

### Feeding behavior

The Automated Recording CAFE, a fly behavior tracking platform, has been described previously^15^. Briefly, one-week-old male adults were transferred into ARC chambers loaded with 300 μL of 1% agar to maintain humidity and provide a water source^39^. Glass capillaries containing liquid food (3.5% sucrose + 3.5% yeast extract, both w/v) were replaced daily at ZT0/CT0, followed by a ∼15-min re-acclimation period. Thus, the effective ARC recording period was ∼23.5 hr per day, which was evenly divided into twelve ∼2-hr food-intake bins. During DD, food was replaced under red light to avoid light stimulation. Flies were typically tracked for >6 days, including 1 day of acclimation, 2 days of LD cycles, and 3 days under constant darkness (DD). Raw data were processed into time series with ∼2-hr intervals, and the analysis of population rhythmicity was performed with easyClock_v3.4^16^ using data from DD1-DD3.

### Feeding/fasting entrainment

Clock[*per*] flies or EEC-labelled Clock[*per*] flies were used for *ad libitum* or feeding/fasting regimens under DD. For feeding/fasting entrainment, flies were provided daily with a solid base diet [5% sucrose, 5% tryptone, and 1% agar (all w/v) supplemented with 0.4% (v/v) propionic acid and 0.06% (v/v) phosphoric acid] for 8 hr starting at CT0, and then fasted on 1% agar for 16 hr. For *ad libitum* controls, flies had continuous access to the same base diet but were transferred to fresh food vials at CT0 and CT8 to match the vial transfers of the experimental group. After 6 cycles of *ad libitum* feeding or feeding/fasting, flies underwent a free-running period prior to sampling. During this period, they were maintained on *ad libitum* feeding for 24 hr (sampling at CT0) or 36 hr (sampling at CT12) under DD. This free-running period was designed to rule out autophagy-related GFP quenching and to test whether clock oscillations driven by feeding/fasting entrainment could be self-sustained in the absence of feeding cues.

### Confocal microscopy

For imaging nuclear GFP or RedStinger (an RFP variant), *Drosophila* midguts were dissected in Dulbecco’s Phosphate-Buffered Saline (D-PBS) and immediately transferred into ice-cold 4% paraformaldehyde (PFA). Once sufficient samples were collected, the cold PFA was replaced with an equal volume of room-temperature 4% PFA, and fixation was carried out for 20 min at room temperature. Following fixation, midguts were washed with D-PBS and mounted using DAPI-containing medium. Midguts with nuclear GFP or RedStinger signals were scanned on a confocal microscope (Olympus FV3000) using 488 nm or 561 nm excitation and 500-540 nm or 570-620 nm detection windows, respectively. For co-localization imaging, line-sequential scanning was used to minimize spectral crosstalk. In preliminary tests, potential GFP channel contamination was excluded by applying a narrower GFP detection range (500-520 nm).

To monitor local clock activity, we used the Clock[*per*] reporter, in which a destabilized nuclear GFP is expressed from a minimal *per* promoter fragment containing CLK/CYC-binding sites^12^. Because the GFP is unstable, Clock[*per*] reflects ongoing CLK/CYC-dependent transcription at the *per* promoter and serves as a time-resolved proxy for molecular clock oscillations in individual midgut cells. To quantify CLK/CYC activity in enterocytes across midgut regions R1-R5, we measured nuclear and cytoplasmic GFP intensities and calculated Δ*F* = *F*_nucleus_ – *F*_cytoplasm_ for each cell, scanning and analyzing each region sequentially in ImageJ^40^. In contrast, due to the low Clock[*per*] signal in midgut EECs, CLK/CYC activity in these cells was evaluated by visually counting Clock[*per*]-positive EECs using a consistent intensity criterion (nuclear GFP clearly above local epithelial background). This strategy provides a semi-quantitative, observer-based estimate of EEC clock activity, rather than an automated, intensity-thresholded measurement.

### Absorption efficiency assay

One-week-old wCS males were randomly divided into 6 groups, with each group further subdivided into vials containing 10 flies each. All flies were acclimated on the base diet (5% sucrose, 5% tryptone, 1% agar, and propionic/phosphoric acids, as described above) for 4 days under LD cycles. Subsequently, flies were transferred to 15 mL conical tubes (10 flies/tube) at 6 evenly distributed Zeitgeber times (1 group/Zeitgeber time). Each tube contained a small food container attached to the cap containing ^35^S-met-labeled food.

Flies were allowed to feed for 2 hr, after which the ^35^S-met-containing caps were replaced with clean caps, and the flies were allowed to excrete for another 2 hr. This period was previously shown to be sufficient for nearly complete excretion under these conditions^41^. Flies and excreta were then collected separately for quantification of ^35^S-met by liquid scintillation counting^42^. For fly bodies, samples were homogenized in 0.2 mL 1× PBS using a motorized pestle prior to counting. Total food intake was estimated from the sum of ^35^S retained in fly bodies and recovered in excreta; absolute nutrient absorption from the ^35^S retained in fly bodies; and absorption efficiency from body ^35^S (absolute absorption) divided by total ^35^S (intake).

### Statistics

Statistical analyses were performed in Python v.3.12.0 unless otherwise specified. Feeding at different times measured by ^35^S-met radiolabeling was modeled as a quadratic function: y = β_0_+ β_1_*x*+ β_2_*x*^2^ + ε, and model coefficients were estimated using ordinary least squares in statsmodels. Spearman correlations between feeding measurements were evaluated using scipy.stats, and multiple comparisons were conducted using ANOVA (Kruskal-Wallis test) followed by Dunn’s post-hoc test. Comparisons between two groups were conducted using the Mann-Whitney U test. Bar and scatter plots are presented as mean ± s.e.m. Circadian rhythms were characterized using Python-JTK based on JTK_CYCLE^43^ and CWT implemented in easyClock (v3.4)^16^, and the corresponding regression modeling and CWT scalograms were generated within easyClock. The easyClock software package is freely available at: https://github.com/HungryFly/easyClock_for_CircadianRhythms.

## Acknowledgments

We thank Dr. Scarlet Park for contributions to the radioisotope-labeling assay, Profs. Mimi Shirasu-Hiza and Phillip Karpowicz for providing *Drosophila* stocks, and Dr. Kathyani Parasram for insightful discussions on enteroendocrine cells. This study was funded by the NIH (R01DC020031).

